# ATM phosphorylates the FATC domain of DNA-PK_cs_ at threonine 4102 to promote non-homologous end joining

**DOI:** 10.1101/2023.02.02.526879

**Authors:** Huiming Lu, Qin Zhang, Daniel J. Laverty, Andrew C. Puncheon, Gareth J. Williams, Zachary D. Nagel, Benjamin PC Chen, Anthony J. Davis

**Affiliations:** Department of Radiation Oncology, UT Southwestern Medical Center, Dallas, Texas, 75390, USA; Department of Environmental Health, Harvard T.H. Chan School of Public Health, Boston, MA, 02115, USA; Department of Biochemistry and Molecular Biology, Robson DNA Science Centre, Charbonneau Cancer Institute, Cumming School of Medicine, University of Calgary, Calgary, AB, Canada

**Keywords:** NHEJ, DNA-PK_cs_, ATM, FATC Domain, Phosphorylation

## Abstract

Ataxia-telangiectasia mutated (ATM) drives the DNA damage response via modulation of multiple signal transduction and DNA repair pathways. Previously, ATM activity was implicated in promoting the non-homologous end joining (NHEJ) pathway to repair a subset of DNA double strand breaks (DSBs), but how ATM performs this function is still unclear. In this study, we identified that ATM phosphorylates the DNA-dependent protein kinase catalytic subunit (DNA-PK_cs_), a core NHEJ factor, at its extreme C-terminus at threonine 4102 (T4102) in response to DSBs. Phosphorylation at T4102 stabilizes the interaction between DNA-PK_cs_ and the Ku-DNA complex and promotes assembly and stabilization of the NHEJ machinery at DSBs. Ablating phosphorylation at this site results in decreased NHEJ, radiosensitivity, and increased radiation-induced genomic instability. Collectively, these findings establish a key role for ATM in NHEJ-dependent repair of DSBs through positive regulation of DNA-PK_cs_.

## INTRODUCTION

DNA double-stranded breaks (DSBs) are cytotoxic DNA lesions that pose an immediate threat to genome stability and failure to properly repair them can lead to cell death, chromosomal aberrations, or carcinogenesis (1). Complex mechanisms collectively termed the DNA damage response (DDR) have evolved in cells to manage DSBs. These mechanisms include DNA damage recognition, activation of signaling cascades and cell cycle checkpoints, chromatin remodeling, transcription regulation, and repair of the DSB. Two members of the phosphatidylinositol-3-kinase-like kinase (PIKK) family, DNA-PK_cs_ (DNA-dependent protein kinase catalytic subunit) and ATM (ataxia telangiectasia-mutated), are instrumental in modulating the cellular response to DSBs (2). DNA-PK_cs_ and ATM are recruited to the site of the DNA damage by the sensors Ku (Ku70 and Ku80 heterodimer) and MRN (MRE11-RAD50-NBS1), respectively, which results in activation of each kinase’s catalytic activity (3). Once activated, DNA-PK_cs_ and ATM phosphorylate substrates that drive multiple pathways responsible for the DDR, including the repair of DSBs.

DNA-PK_cs_ promotes DSB repair via the non-homologous end joining (NHEJ) pathway (4). In mammalian cells, NHEJ is the pathway primarily responsible for the repair of radiation-induced DSBs and DSBs generated for V(D)J and class switch recombination (CSR) during T- and B-cell lymphocyte maturation. NHEJ initiates when Ku rapidly binds directly to the DSB, where it then performs its primary function as a scaffold to recruit the NHEJ machinery to the damage site. It is still unclear if the recruitment of the NHEJ factors occurs in a stepwise sequential manner or via a dynamic assembly, but early studies found that the core NHEJ factors, Ku, DNA-PK_cs_, X-Ray Repair Cross Complementing 4 (XRCC4), DNA Ligase 4 (LIG4), and XRCC4-like factor (XLF) collectively stabilize at the DNA damage site via a number of protein-protein interactions and are modulated by DNA-PK_cs_ kinase activity (5). Single-molecule FRET studies revealed that NHEJ forms two synaptic complexes, referred to as “long-range” and “short-range” (6). The long-range complex synapses two DSB ends but they are more than 100 Å apart, whereas the DNA ends are more closely aligned in the short-range synaptic complex. DNA-PK_cs_ kinase activity was found to be important for the transition from the long-range to the short-range complex. Single-particle cryo-electron microscopy recently offered insight into the factors that may mediate the NHEJ synaptic complexes (7). The long-range complex contained the Ku heterodimer, DNA-PK_cs_, LIG4, XRCC4, and XLF. In the absence of DNA-PK_cs_, Ku, LIG4, XRCC4, and XLF formed a complex in which the DNA ends are horizontal and aligned for processing and ligation, similar to what is predicted for the short-range synaptic complex. Furthermore, evidence suggests that DNA-PK_cs_ activity promotes the processing of unligatable DNA ends by regulating the recruitment and stabilization of DNA end processing factors at DSBs and via autophosphorylation-induced conformational changes that free the DSB ends (4,8–10). However, there is still significant debate about the role and function of DNA-PK_cs_, DNA-PK_cs_ activity, and DNA-PK_cs_ phosphorylation status in synaptic complex formation and transition (11).

ATM, the protein defective in the heritable disorder ataxia telangiectasia (A-T), is the kinase primarily responsible for signal transduction and cell cycle regulation in response to DSBs (12). Cells derived from A-T patients are hypersensitive to DSB generating agents, suggesting that ATM is essential for the repair of DSBs. ATM does not play a direct role in DSB repair but influences multiple pathways via phosphorylation events. First, ATM promotes homologous recombination (HR) by phosphorylating multiple HR factors, including MRE11, NBS1, PALB2, BRCA1, and CtIP (13–17). Second, ATM is required for NHEJ at complex lesions, DSBs induced in heterochromatin, and DSBs harboring a blocked DNA end (18,19). Moreover, ATM phosphorylates many components of the NHEJ pathway, such as XLF, Artemis, and DNA-PK_cs_, but there is limited knowledge of the functionality of these phosphorylation events (20). ATM plays at least one important additional role in NHEJ that has not been elucidated, but it is redundant with the core NHEJ factor XLF and thus not normally apparent (21). Finally, it has been reported that ATM is dispensable for NHEJ but promotes repair fidelity via an undefined mechanism (22).

Here, we uncover a novel function for ATM in NHEJ via phosphorylating DNA-PK_cs_. The data show that DNA-PK_cs_ is phosphorylated by ATM in its FATC domain at threonine 4102 in response to DSBs. Phosphorylation at T4102 promotes stabilization of the interaction between DNA-PK_cs_ and the Ku-DNA complex. Ablating phosphorylation of DNA-PK_cs_ at threonine 4102 results in destabilization of the NHEJ machinery at DSBs, leading to decreased NHEJ efficiency and genomic instability. Our results establish a model in which phosphorylation of the FATC domain of DNA-PK_cs_ by ATM promotes efficient NHEJ.

## Material & Methods

### Cell lines, cell culture, inhibitor treatments, and cell cycle synchronization

Chinese Hamster Ovary (CHO) V3 cells and V3 cells stably expressing DNA-PK_cs_ wild-type, T4102A, and T4102D were cultured in Hyclone α-minimum Eagle’s medium supplemented with 5% newborn calf serum, 5% fetal bovine serum (Corning), and 1X penicillin/streptomycin (Gibco). U2OS cells were cultured in Dulbecco’s Modified Eagle Medium supplemented with 10% fetal bovine serum and 1X penicillin/streptomycin (Gibco). All cells were grown in an atmosphere of 5% CO_2_ at 37°C. Ionizing radiation by γ-rays was achieved by incubating cells in Mark 1 ^137^Cs irradiator (JL Shepherd and Associates) at room temperature, for a total dose of 10 Gy unless otherwise indicated. To inhibit DNA-PK_cs_, ATM, or ATR, cells were incubated for 2 hr prior to irradiation with 5 μM NU7441 (SelleckChem), 5 μM KU55933 (SelleckChem) or 5 μM VE821 (SelleckChem), respectively. Enrichment of U2OS cells at G1 phase and S/G2 were performed via a double thymidine block as previously described (23).

### Irradiation

Cells were irradiated with γ-rays generated by a Mark 1 ^137^Cs irradiator (J.L. Shepherd and Associates) for 10 Gy, or at the doses denoted in the figures.

### Generation of antibody against phosphorylated DNA-PK_cs_ at T4102

Anti-pT4102 polyclonal antibodies were generated by immunizing New Zealand white rabbits with KLH (keyhole limpet hemacyanin)-conjugated phosphopeptide GLSEET[PO_3_]QVKCLMC. The phospho-specific antibodies were first passed through the corresponding unphosphorylated peptide-conjugated Sepharose CL-4B column (Pierce) to deplete IgGs that were not phospho-specific. The flow through IgGs were then affinity-purified using a phospho-peptide column. Eluted anti-pT4102 polyclonal antibodies were verified with phospho and non-phospho peptides via dot plot analysis.

### Site-directed point mutagenesis

Site-directed mutagenesis was performed using PCR to substitute T4102 on DNA-PK_cs_ with alanine or aspartic acid using the pCMV-F2-K and pCMV-YFP-K as a template to generate mutated FLAG-tagged and YFP-tagged DNA-PK_cs_, respectively. The primers are used were: T4102A: T4102A Forward 5’-TGGGCTTTCAGAAGAGGCTCAAGTGAAGTGCCT-3’ and T4102A Reverse 5’-AGGCACTTCACTTGAGCCTCTTCTGAAAGCCCA-3’ and T4102D: T4102D Forward 5’-GTGGGCTTTCAGAAGAGGATCAAGTGAAGTGCCTG-3’ and T4102D Reverse 5’-CAGGCACTTCACTTGATCCTCTTCTGAAAGCCCAC-3’.

### Immunoblotting

Immunoblotting was performed as previously described (24). The following antibodies were used in this study: antibodies from anti-DNA-PK_cs_ phospho-T4102 (made in this study as described above), anti-DNA-PK_cs_ phospho-S2056 (Abcam, ab124918), anti-ATM phospho-S1981 (Abcam, ab81292), anti-CHK2 phospho-T68 (Cell Signaling, 2197), anti-LIG4 (Cell Signaling, 14649), anti-Ku80 (Santa Cruz, sc-17789), anti-Ku70 (Santa Cruz, sc-515736), anti-XRCC4 (Santa Cruz, sc-271087), anti-XLF (Santa Cruz, sc-166488), anti-GFP (Santa Cruz, sc-8334), anti-KAP1 (Bethyl Laboratories A300-274A), anti-KAP1 phospho-S824 (Bethyl Laboratories, A300-767A), anti-tubulin (Sigma-Aldrich, T5168), anti-FLAG M2 (Sigma-Aldrich, F1804), anti-phospho-H2AX (S139) (EMD Millipore, 05-636), and anti-Histone H3 antibody (Biolegend, 819411). Mouse monoclonal antibodies against DNA-PK_cs_ (Clone # 25-4) were produced in house. Secondary antibodies used include anti-mouse IgG (HRP-linked) (Cell Signaling, 7076) and anti-rabbit IgG (HRP-linked) (Cell Signaling, 7074).

### Immunoprecipitation (IP) Assays

To detect phosphorylation of DNA-PK_cs_ at T4102, V3 cells stably expressing YFP-tagged wild-type DNA-PK_cs_ were irradiated with a dose of 10 Gy and allowed to recover for 30 min. The cells were then washed twice with cold PBS, harvested, and lysed using IP lysis buffer (50 mM Tris-HCl pH 7.4, 500 mM NaCl, 0.5% NP-40, and 10% Glycerol with 1X protease and phosphatase inhibitor cocktails (Thermo Fisher)). The lysates were sonicated on ice and then cleared of cellular debris by centrifuging at 20,000 × g for 30 min. 2 mg of total protein was incubated with the DNA-PK_cs_ monoclonal antibody (25–4) and Protein A/G beads (Thermo Fisher) overnight at 4° C with mixing. The following day the beads were washed three times with lysis buffer and then twice with washing buffer (50 mM Tris-HCl pH 7.4, 150 mM NaCl, 0.5% NP-40, and 10% Glycerol). Following the final wash, the beads were divided two parts, with one resuspended in 1X SDS sample buffer and other treated with lambda protein phosphatase (λPP) (New England Biosciences) at 30° C for 30 min to remove phosphorylation events, the samples were washed, and finally resuspended in 1X SDS sample buffer. The samples were resolved via SDS-PAGE and phosphorylation at T4102 was assessed via immunoblotting.

To investigate protein-protein interactions, CHO V3 cells and V3 cells complemented with WT, T4102A, or T4102D were irradiated with a dose of 10 Gy and allowed to recover for 10 min. Subsequently, the irradiated cells were washed three times in cold PBS, harvested, and lysed using immunoprecipitation (IP) Lysis buffer (50 mM Tris-HCl pH 7.4, 150 mM NaCl, 2 mM MgCl_2_, 0.4% NP-40, 0.6% Triton X-100, 1X protease inhibitor cocktail, 1X phosphatase inhibitor cocktail 2, 20 U/mL Benzonase (Novagen)) on ice. The lysates were sonicated on ice and then cleared of cellular debris by centrifuging at 20,000 × g for 30 min. 2 mg of total protein was incubated with 2 μg anti-GFP antibody, and 30 μL of Protein A/G magnetic agarose beads (Thermo Fisher) overnight with spinning at 4° C. The beads were washed with IP washing buffer (20 mM Tris-HCl pH 7.4, 150 mM NaCl, 0.2% Triton X-100) for 5 times, and boiled in 1X SDS sample buffer. The samples were resolved via SDS-PAGE and immunoblotting was performed for the proteins indicated in the figures.

### In vitro phosphorylation assay

Purification of ATM from HT1080 cells stably expressing FLAG-tagged ATM and DNA-PK_cs_ from CHO V3 cells stably expressing YFP-tagged DNA-PK_cs_ was performed as previously described with some modifications (23). To activate the ATM protein, the cells were irradiated with a dose of 10 Gy and allowed to recover for 30 min. Next, the cells were washed three times with cold PBS, harvested, and lysed using Purification Lysis Buffer (50 mM Tris-HCl pH 7.4, 500 mM NaCl, 0.5% NP-40, and 10% Glycerol with 1X protease and phosphatase inhibitor cocktails (Thermo Fisher)). The lysates were sonicated on ice and then cleared of cellular debris by centrifuging at 20,000 × g for 30 min. 2 mg of total proteins was incubated with 30 μL M2 FLAG magnetic beads (Sigma-Aldrich) for ATM and 2 mg DNA-PK_cs_ antibody and 30 μL Protein A/G beads for DNA-PK_cs_. After an overnight incubation at 4° C, the beads were washed twice with Purification Lysis Buffer, twice with Wash Buffer 1 (50 mM Tris-HCl pH 7.4, 150 mM NaCl, 0.5% NP-40, and 10% Glycerol) and finally twice with Wash Buffer 2 (20 mM Tris-HCl pH 7.4, 50 mM KCl, and 10% Glycerol). For ATM protein, half of beads were resuspended in 1X SDS sample buffer and the sample was resolved via SDS-PAGE, and the other half was for the in vitro phosphorylation assay. To remove phosphorylations associated with the purified DNA-PK_cs_ protein, the beads were further equilibrated in 1X λPP reaction buffer (New England Biosciences) and treated with λPP at room temperature for 30 min. The sample was washed using the protocol described above. One-third of the beads containing DNA-PK_cs_ was resolved via SDS-PAGE, and the rest were utilized for the ATM-mediated in vitro phosphorylation assay. The SDS-PAGE gel that resolved purified YFP-DNA-PK_cs_ and FLAG-ATM was Coomaissie Blue stained to show purification of each protein.

In vitro phosphorylation of DNA-PK_cs_ by ATM was conducted by mixing beads containing YFP-DNA-PK_cs_ with beads containing FLAG-tagged ATM or control beads. The bead mixtures were placed in Kinase Reaction Buffer (20 mM HEPES 7.4, 50mM KCl, 2 mM MgCl2, 2mM ATP, 1mM DTT and 5% Glycerol) and incubated at 37° C for 1 hr with mixing. The reactions were terminated by adding 1X SDS sample buffer and boiling the samples at 95° C for 5 min. The phosphorylation of DNA-PK_cs_ at T4102 was then assessed by immunoblotting with the pT4102 antibody.

### Fluorescent Immunostaining and microscopy

IR-induced 53BP1 foci kinetics were monitored in G1 cells as previously described with modifications (25,26). Briefly, CHO V3 cells and V3 cells complemented with WT, T4102A, or T4102D were seeded on “PTFE” Printed Slides (Electron Microscopy Sciences) and two days later the cells were mock treated or irradiate with a dose of 2 Gy. At different time points after IR (0.5, 1, 3 or 7 hours), the cells were washed twice with cold PBS and fixed with 4% paraformaldehyde (in PBS) for 20 min at room temperature, washed five times with PBS, and incubated in 0.5% Triton X-100 on ice for 10 min. Cells were washed five times with 1× PBS and incubated in blocking solution (5% goat serum (Jackson Immuno Research) in 1× PBS) for 1h. The blocking solution was replaced with the 53BP1 (ab175933, Abcam) and Cyclin A2 (ab16726, Abcam) primary antibodies (1:1000 dilution for both antibodies) diluted in 5% normal goat serum in 1× PBS and the cells were incubated at 4° C overnight. The next day the cells were washed five times with Wash Buffer (1% BSA in 1× PBS). Next, the cells were incubated with anti-rabbit IgG conjugated with Alexa Fluor 488 (Molecular Probes) and anti-mouse IgG conjugated with Texas Red (Molecular Probes) (1:1000 dilution for both antibodies) secondary antibodies in 1% BSA, 2.5% goat serum in 1× PBS for 1 h in the dark, followed by five washes. After the last wash, the cells were mounted in VectaShield Antifade mounting medium containing 4′,6-diamidino-2-phenylindole (DAPI). Images were acquired using a Zeiss AxioImager fluorescence microscope utilizing a 63× oil objective lens. The 53BP1 foci were only counted in the cells with no Cyclin A staining.

### Laser micro-irradiation and real-time recruitment

Real-time recruitment of fluorescent tagged DNA-PK_cs_ WT, T4102A, T4102D, Ku80, XRCC4, XLF, and PKNP in response to DSB induction was examined following laser micro-irradiation with a Carl Zeiss Axiovert 200M microscope with a Plan-Apochromat 63×/NA 1.40 oil immersion objective (Carl Zeiss) as previously described (24,26). CHO V3 cells complemented with YFP-tagged WT, T4102A, or T4102D or GFP-tagged Ku80, XRCC4, XLF, or PNKP was transiently expressed in CHO V3 cells complemented with FLAG-tagged WT, T4102A, or T4102D were seeded on a 35 mm glass-bottomed dish (Mattek) and incubated with 10 μM BrdU. 24 h later, the medium was replaced with CO_2_-independent medium and placed in a chamber on the microscope that was set at 37° C. To generate laser-induced DSBs, a 365-nm pulsed nitrogen laser (Spectra-Physics, Catalog #VSL337NDS2, purchased in May 2020) was set at 80% of maximum power output and micro-irradiation was performed using the pulsed nitrogen laser. Time-lapse images were taken using an AxioCam HRm camera (Carl Zeiss). Carl Zeiss Axiovision software (v4.91) was used to measure fluorescence intensities of the micro-irradiated and control areas, and the resulting intensity of irradiated area was normalized to non-irradiated control area to obtain the alteration of the interested proteins as described previously (24,26).

### Subcellular fractionation

The accumulation of DNA damage response proteins to chromatin following IR-induced DNA damage was examined using the Subcellular Protein Fractionation Kit (Thermo Fisher) as previously described (23). Briefly, the V3 cells expressing either DNA-PK_cs_ WT or T4102A were mock-treated or irradiated with 10 Gy, allowed to recover for 10 min, harvested after trypsinization, and then processed with the Thermo Fisher Subcellular Protein Fractionation Kit according to the manufacturer’s instructions. The protein concentration of each sample was measured using a Pierce BCA Protein Assay kit (Thermo Fisher). 20 μg of each fraction was resolved via SDS-PAGE, and then transferred to a PVDF membrane for immunoblotting

### NHEJ assay

CHO cells were seeded in triplicate at a density of 40,000 cells per well in 12-well plates. The following day, cells were transfected with reporter plasmids using Lipofectamine 3000 (ThermoFisher) according to the manufacturer protocol. For each cell line, one well was transfected with undamaged plasmid cocktail, one well was transfected with NHEJ plasmid cocktail, and one well was left untransfected. After 24 hr, cells were trypsinized and analyzed by flow cytometry using an Attune NxT flow cytometer. Compensation and gating were established by running untransfected cells along with cells transfected with individual undamaged plasmids expressing wild type fluorescent proteins (single color controls) as described previously (27). Each experiment was conducted three times on separate days.

Reporter plasmid cocktails were as follows: undamaged plasmid cocktail: 50 ng pMax_BFP, 50 ng pMax_mOrange, 500 ng non-fluorescent carrier plasmid pDDCMV. NHEJ plasmid cocktail: 50 ng BFP_NHEJ, 50 ng pMax_mOrange, 500 ng non-fluorescent carrier plasmid pDDCMV. Reporter fluorescence was used as a measurement of NHEJ efficiency as described in Piett et. al. and below:

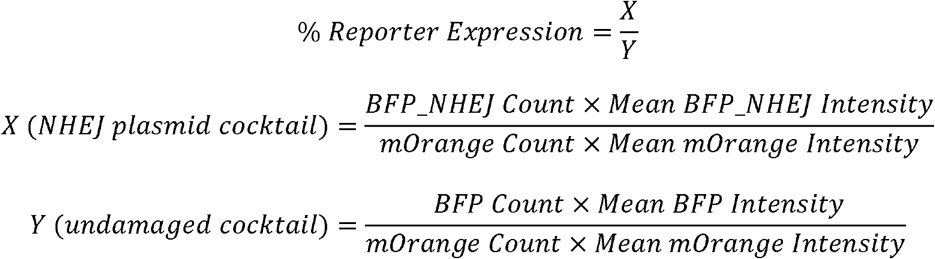

### Colony formation assay

Cell survival curves were obtained by measuring the colony-forming abilities of irradiated cell populations as previously described (28). CHO V3 cells and V3 cells complemented with WT, T4102A, or T4102D were mock treated or irradiated at doses of 1, 2, 4, or 6 Gy and then plated on 60-mm plastic Petri dishes. After 10 days, cells were fixed with 100% ethanol and stained with 0.1% crystal violet in a 100% ethanol solution. Colonies were scored and the mean value for triplicate culture dishes was determined. Cell survival was normalized to plating efficiency of untreated controls for each cell type.

### Chromosome aberration assay

To investigate genome stability following irradiation in CHO V3 cells and V3 cells complemented with WT, T4102A, or T4102D, the cells were irradiated with a dose of 2 Gy and then allowed to recover in normal cell culture conditions for 24 hrs. Next, the cells were incubated with Colcemid at concentration of 0.1 ug/mL for 4 hrs and then harvested by trypsinzation. After washing with PBS at room temperature, the cells were then incubated with warmed (37°C) 75 mM KCl. Samples were processed and chromosomal abnormalities were scored as previously described (25).

## Results

### ATM phosphorylates DNA-PK_cs_ in its FATC domain at threonine 4102 in response to DNA double strand breaks

DNA-PK_cs_ is composed of HEAT (Huntington-elongation factor 3A- PP2A subunit-TOR) repeats, which are differentiated into a unique N-terminal domain (N-HEAT, amino acids 1-892) and a central unit called the M-HEAT domain (amino acids 893-2801) and a C-terminal region that contains the kinase domain (amino acids 3565-4100), which is flanked N-terminally by the FAT (FRAP, ATM, TRRAP) domain and C-terminally by the FATC (FAT C-terminal) domain (29). The FATC domain comprises the extreme C-terminus of DNA-PK_cs_ and is a highly conserved domain of approximately 30 amino acids (30). Early work showed that the FATC domain is indispensable for DNA-PK_cs_ activity, as deletion or mutagenesis in this domain deleteriously affects the function of DNA-PK_cs_ (31). This drove us to postulate that modulation of DNA-PK_cs_ may be regulated by a post-translation modification in the FATC domain. Previous work identified that DNA-PK_cs_ is phosphorylated in the FATC domain at threonine T4102 (pT4102)(32), however the biological relevance of this phosphorylation event has been not previously investigated, propelling us to determine if this site regulates the function of DNA-PK_cs_. Alignment of DNA-PK_cs_ orthologues identified that T4102 is conserved in nearly all vertebrates, which further provided evidence that phosphorylation at this site may be an evolutionary conserved regulatory mechanism (Fig. 1A and Sup. Fig. 1). Analysis of the DNA-PK_cs_ structure shows that T4102 is surface exposed and thus a viable phosphorylation target (Fig. 1B)(33). To examine if DNA-PK_cs_ is phosphorylated at this amino acid after DNA damage in cells, a phospho-specific antibody to T4102 was generated. DNA-PK_cs_ is phosphorylated at T4102 (pT4012) following exposure to ionizing radiation (IR) and the signal was lost when the sample was treated with lambda phosphatase, supporting that the antibody recognizes a phosphorylation event (Fig. 1C). Next, the specificity of the phospho-antibody was assessed by complementing the DNA-PK_cs_ deficient Chinese Hamster Ovary (CHO) cell line V3 with wild-type DNA-PK_cs_ (WT) or DNA-PK_cs_ in which the phosphorylation site at T4102 was ablated via alanine substitution (T4102A) (Sup. Fig. 2A). The signal was significantly decreased in cells expressing the T4102A mutant protein, indicating the antibody specifically recognizes pT4102 (Fig. 1D). Furthermore, pT4102 occurs following IR exposure in a dosage-dependent manner (Fig. 1E), with observation of the pT4102 signal starting at 5 minutes, peaking at 30-60 minutes, and still detectable 24 hours post-IR (Fig. 1F). This phenotype is conserved in human cells, as pT4102 initiates 5 minutes and peaks 60 minutes post-IR in the human cell line U2OS (Sup. Fig. 2B) and IR-induced pT4102 is also observed in HeLa cells (Sup. Fig. 2C). In addition to IR, treatment with the radiomimetic agent neocarzinostatin (NCS) and the topoisomerase 2 inhibitor etoposide (ETO) induced robust pT4102 signal (Fig. 1G). Modest phosphorylation of DNA-PK_cs_ at T4102 was observed after treatment with the topoisomerase 1 inhibitor camptothecin (CPT), but limited to no phosphorylation occurred following treatment with the DNA cross-linking agent mitomycin (MMC) and DNA alkylating agent methyl methanesulfonate (MMS) (Fig. 1G). As IR, NCS, and ETO directly generate DSBs and CPT-induced DNA damage can be processed to form DSBs, the data suggests that DSBs induce phosphorylation of DNA-PK_cs_ at T4102. We then assessed if IR-induced phosphorylation of DNA-PK_cs_ at T4102 is cell cycle specific. Cells were synchronized via a double thymidine block and then released to allow examination of IR-induced phosphorylation of DNA-PK_cs_ at T4102 in G1 and S/G2 phases of the cell cycle. We observed that pT4102 occurs in both G1 and S/G2 phases, but phosphorylation is more prominent in G1 phase of the cell cycle (Fig. 1H). We next aimed to identify the kinase responsible for phosphorylating DNA-PK_cs_ at T4102 in response to DNA damage. As phosphorylation at this site is induced by DSBs, we focused on the DNA damage-responsive kinases, including DNA-PK_cs_, ATM, and ATR. Pretreatment of cells with the inhibitors NU7441, KU55933, and VE-821 to block DNA-PK_cs_, ATM, and ATR activity, respectively, shows that pT4102 is significantly attenuated in cells pretreated with KU55933 but not NU7441 or VE-821 (Fig. 1I). This data indicates that ATM phosphorylates DNA-PK_cs_ at T4102 in response to DNA damage. To verify this result, we assessed pT4102 in the ATM deficient cell line AT5 and AT5 cells complemented with ATM (34). IR-induced phosphorylation of DNA-PK_cs_ at T4102 is almost completely lost in the ATM-deficient cell line and this was rescued by complementing AT5 cells with wild-type ATM (Fig. 1K). Finally, we examined the ability of ATM to phosphorylate DNA-PK_cs_ at T4102 *in vitro*. DNA-PK_cs_ was isolated from cells and incubated with purified ATM in the presence of ATP (Sup. Fig. 3). ATM autophosphorylates at S1981 in this system and it phosphorylates DNA-PK_cs_ at threonine 4102 as monitored by the pT4102 antibody (Fig. 1J). Collectively, the data illustrates that ATM phosphorylates DNA-PK_cs_ at T4012 in response to DNA damage.

**Figure 1.**
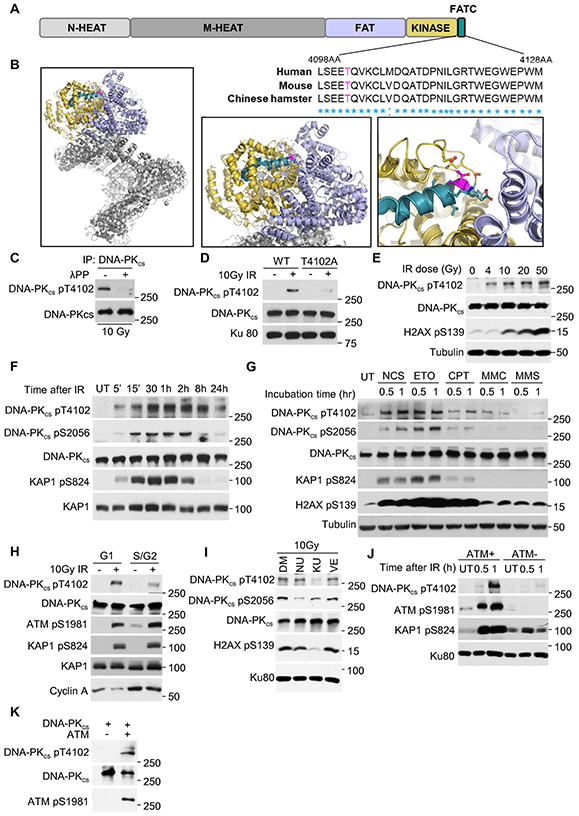
ATM phosphorylates DNA-PK_cs_ in its FATC domain at threonine 4102 in response to DSBs. **(A)** Threonine 4102 (T4102) is conversed in the FATC domain of DNA-PK_cs_ in human, mouse, and Chinese hamster. **(B)** Structure of DNA-PK_cs_, generated in PyMol version 2.5 from PDB: 7OTP (33). T4102 residue is highlighted in magenta. DNA-PK_cs_ domains are highlighted thusly: FATC domain (green), kinase domain (yellow), FAT domain (blue), and N-terminal HEAT repeat domains (gray). **(C)** DNA-PK_cs_ is phosphorylated at T4102 (pT4102) after ionizing radiation (IR) and this signal is lost when treated with lambda protein phosphatase (λPP). (**D)** Ablating the T4102 phosphorylation site via alanine substitution (T4102A) results in loss of IR-induced phosphorylation at T4102. **(E)** Tracking IR-induced dose-dependent pT4102. **(F)** Time course of pT4102 following treatment with 10 Gy of IR. **(G)** Phosphorylation of DNA-PK_cs_ at T4102 following treatment with various DNA damaging agents for 1 hour, including 200 ng/mL neocarzinostatin (NCS), 1 μM etoposide (ETO), 1 μM camptothecin (CPT), 0.5 μg/mL mitomycin C (MMC), and 50 μg/mL methyl methanesulfonate (MMS). **(H)** Cell cycle-dependent phosphorylation of DNA-PK_cs_ at T4102 following IR. **(I)** Inhibition of ATM, but not ATR or DNA-PK_cs_, blocks IR-induced DNA-PK_cs_ phosphorylation at T4102. Inhibitors include DNA-PK_cs_ inhibitor NU7441 (NU), ATM inhibitor KU55933 (KU), and ATR inhibitor VE821 (VE). **(J)** IR-induced phosphorylation of DNA-PK_cs_ at T4102 is lost in the ATM-deficient cell line AT5BIVA. **(K)** ATM phosphorylates DNA-PK_cs_ *in vitro*.

### Phosphorylation at T4102 stabilizes the interaction between DNA-PK_cs_ and the Ku-DNA complex

As T4102 lies in the FATC domain, we next assessed if phosphorylation of this site modulates DNA-PK_cs_ kinase activity. We complemented V3 cells with YFP-tagged WT, T4102A, and the DNA-PK_cs_ mutation in which T4102 was mutated to aspartic acid to mimic phosphorylation at this amino acid (T4102D) (Sup. Fig. 2A) and subsequently examined DNA-PK_cs_-mediated phosphorylation in response to DSBs. IR-induced autophosphorylation of DNA-PK_cs_ at S2056 is decreased in T4102A cells compared to WT cells (Fig. 2A). Furthermore, phosphorylation of KAP1 at serine 824 and phosphorylation of H2AX at serine 139 are decreased in T4102A cells at early time points post-IR treatment (Fig. 2A), which is consistent with our previous data showing that DNA-PK_cs_ phosphorylates these sites (24). Next, we determined if the decrease in DNA-PK_cs_-mediated phosphorylation events was due to a decrease in the DNA-PK_cs_-Ku interaction. The data shows that T4102A has decreased interaction with Ku80 and Ku70 following exposure to IR, compared to WT and T4102D (Fig. 2B). The disruption of the DNA-PK_cs_-Ku interaction was further examined via monitoring the dynamics of DNA-PK_cs_ at laser-generated DSBs. We found that blocking T4102 phosphorylation results in decreased recruitment of DNA-PK_cs_ to DSBs compared to WT and the T4102D proteins (Fig. 2C). Moreover, kinetic analysis shows that the T4102A mutant prematurely dissociates from laser-generated DSBs compared to the WT and T4102D proteins (Fig. 2D). Collectively, the data illustrates that blocking DNA-PK_cs_ phosphorylation at T4102 destabilizes the DNA-PK_cs_-Ku complex, resulting in decreased DNA-PK_cs_-mediated phosphorylation events following DSB induction.

**Figure 2.**
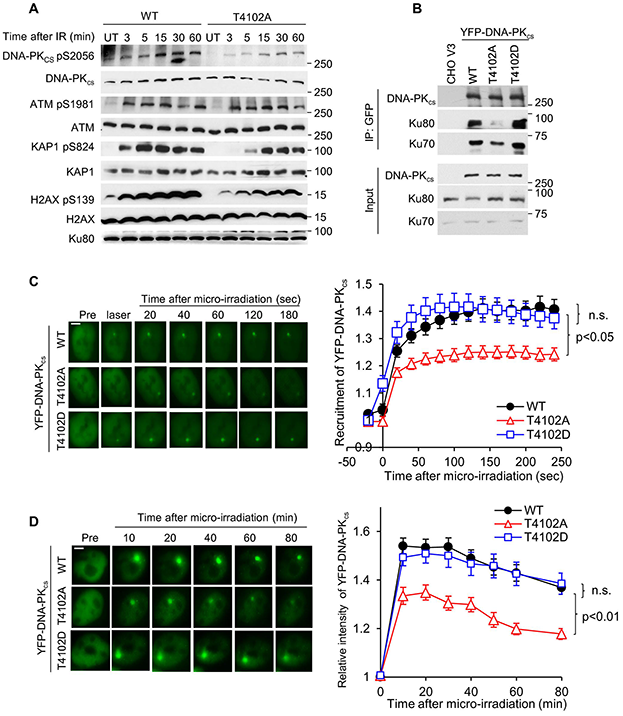
Phosphorylation at T4102 stabilizes the interaction between DNA-PK_cs_ and the Ku-DNA complex. **(A)** Blocking DNA-PK_cs_ phosphorylation at T4102 suppresses IR-induced DNA-PK_cs_-mediated phosphorylation events. **(B)** T4102 phosphorylation promotes stabilization of the interaction between DNA-PK_cs_ and the Ku70/80 heterodimer after IR. **(C)** Ablating T4102 phosphorylation attenuates the initial recruitment of DNA-PK_cs_ to laser-induced DSBs. Relative fluorescent intensity of YFP-tagged wild type DNA-PK_cs_ WT, T4102A, and T4102D are presented as mean ± SEM and significance was assessed via a student’s t-test. The bar indicates 5 μm. **(D)** T4102 phosphorylation promotes accumulation of DNA-PK_cs_ at laser-induced DSBs. Relative fluorescent intensity of YFP-tagged wild type DNA-PK_cs_ WT, T4102A, and T4102D are presented as mean ± SEM and significance was assessed via a student’s t-test. The bar indicates 5 μm.

### Stabilization of the NHEJ machinery at DSBs is promoted by phosphorylation of DNA-PK_cs_ at T4102

As DNA-PK_cs_ has been implicated in promoting the long-range synaptic complex and T4102 promotes stabilization of the DNA-PK_cs_-Ku complex, we next assessed if phosphorylation at this site affects the dynamics of the NHEJ machinery at DSBs. First, the recruitment of the core NHEJ factors Ku80, XRCC4, and XLF to laser-generated DSBs was examined in V3 cells complemented with FLAG-tagged DNA-PK_cs_ WT, T4102A, and T4102D (Sup. Fig. 4A). The initial dynamics of Ku80 is similar in all three cell lines, illustrating that T4102 phosphorylation does not regulate the ability of the Ku heterodimer to bind to DSBs or its initial dynamics at DSBs (Sup. Fig. 4B). We observed that GFP-tagged XRCC4 and XLF are quickly recruited to laser-induced DSBs in WT, T4102A, and T4102D cells (Fig. 3A & B). However, XRCC4 and XLF signal wanes in the T4102A complemented cells compared to the WT and T4102D cells (Fig. 3A & B), suggesting that stability of the NHEJ machinery is decreased when phosphorylation of DNA-PK_cs_ at T4102 is blocked. Moreover, we extended this to the DNA end processing factor PNKP. Similar to XRCC4 and XLF, PNKP recruitment/retention at laser-generated DSBs was decreased in T4102 complemented cells compared to WT and T4102D (Fig. 3C). Finally, we observed that IR-induced recruitment of XRCC4 and LIG4 to the chromatin faction was significantly decreased in the T4102A cells compared to WT cells, supporting the observation that blocking T4102 phosphorylation affects the dynamics of NHEJ factors at DSBs (Sup. Fig. 5). Together, the data show that ATM-mediated phosphorylation of DNA-PK_cs_ at T4102 promotes stabilization of the NHEJ machinery at DSBs.

**Figure 3.**
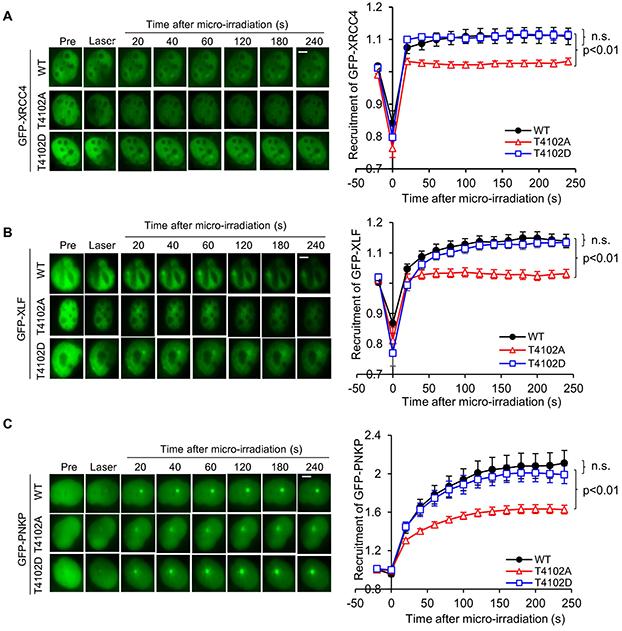
Stabilization of the NHEJ machinery at DSBs is promoted by phosphorylation of DNA-PK_cs_ at T4102. Ablating T4102 phosphorylation attenuates the recruitment of GFP-XRCC4 (**A**), GFP-XLF (**B**), and GFP-PNKP (**C**) to laser-generated DSBs. Relative fluorescent intensity of GFP-tagged XRCC4 (A), XLF (B), and PNKP (C) in CHO V3 cells with FLAG-tagged DNA-PK_cs_ WT, 4102A, and T4102D are presented as mean ± SEM and significance was assessed via a student’s t-test. The bar indicates 5 μm.

### Phosphorylation of DNA-PK_cs_ at T4102 is important for NHEJ-mediated DSB repair

As blocking phosphorylation of DNA-PK_cs_ at T4102 destabilizes the NHEJ machinery at DSBs, we next examined if blocking phosphorylation at this site affects NHEJ. First, we assessed IR-induced 53BP1 focus formation and resolution in G1 cells, which was used as an indirect marker for DSB repair and NHEJ. As shown in Fig. 4A, we found at 30 minutes post-IR, the number of 53BP1 foci is the same in V3, WT, T4102A, and T4102D cells. However, at 1-, 3-, and 7-hours post-IR, 53BP1 focus resolution was attenuated in V3 and T4102A cells compared to WT and T410D cells. 53BP1 focus resolution is more attenuated in the DNA-PK_cs_-deficient V3 cells compared to the T4102A cells, suggesting that T4102 phosphorylation does not result in a complete abolishment of NHEJ. Next, NHEJ was monitored via a FM-HCR assay (27). We observed that the loss of DNA-PK_cs_ (V3) results in almost complete loss of NHEJ, which was rescued by expression of WT DNA-PK_cs_ (Fig. 4B). The T4102A mutant complemented cells show a modest, but significant decrease in NHEJ compared to WT. The T4102D complemented cells show a modest, but significant increase in NHEJ efficiency compared to WT cells, indicating that phosphorylation of T4102 promotes NHEJ. As NHEJ is decreased in T4101A cells, we determined if blocking this phosphorylation site results in increased sensitivity to radiation. As shown in Fig. 4C, T4102A cells are modest radiosensitive compared to WT and T4102D cells. Finally, IR-generated chromosomal aberrations, as assessed by chromatin and chromosome breaks and chromosome fusions, are increased in T4102A cells compared to WT and T4102D cells (Fig. 4D & E). Collectively, the findings in this study show that ATM phosphorylates the FATC domain of DNA-PK_cs_ at T4102 in response to DSBs. Moreover, phosphorylation at T4102 promotes stabilization of the NHEJ machinery at DSBs and ultimately NHEJ and genome stability. This study provides further insight into how ATM functions in the response to DSBs by promoting NHEJ.

**Figure 4.**
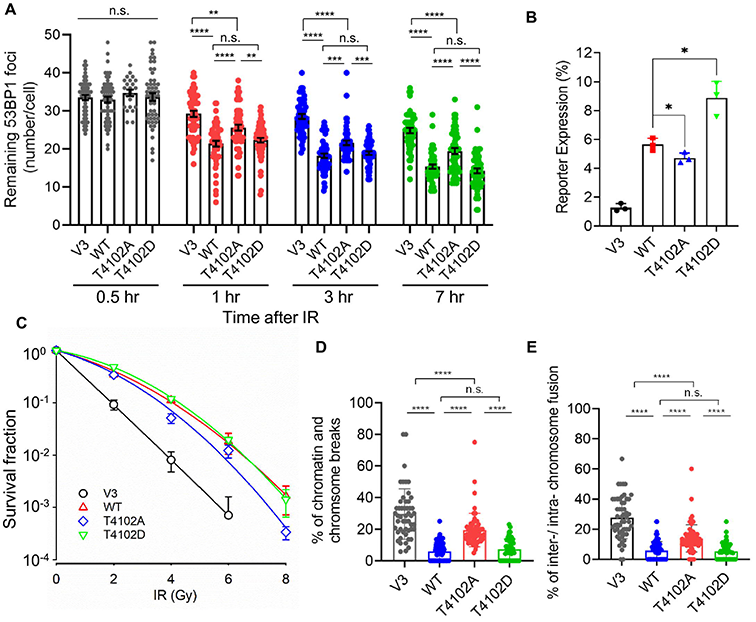
Phosphorylation of DNA-PK_cs_ at T4102 is important for NHEJ-mediated DSB repair. **(A)** IR-induced 53BP1 foci resolution in G1 cells is attenuated in T4102A cells compared to WT and T4102D cells. CHO V3 cells and V3 cells complemented with WT, T4102A, or T4102D were irradiated with 2 Gy of γ-rays and 53BP1 foci formation and resolution was assessed 0.5, 1, 3 and 7 h post-IR. 53BP1 foci at each time point were calculated in over 50 Cyclin A-negative cells and the data are presented as Mean ± SEM. Student’s t-test (two-sided) was performed to assess statistical significance (n.s., not significant; **, p<0.01; ***, p<0.001; ****, p<0.0001). **(B)** DNA-PK_cs_ phosphorylation at T4102 promotes NHEJ. NHEJ reporter and control plasmids were transfected into CHO V3 cells and V3 cells stably expressing YFP-tagged DNA-PK_cs_ WT, T4102, and T4102D and NHEJ efficiency was calculated using flow cytometry analysis as described in the Materials and Methods. The data are presented as mean±SD with p-value from three independent repeats. *, p<0.05. **(C)** T4102A complemented cells are radiosensitive compared to WT and T4102D cells. Clonogenic survival assays were performed to compare the radiation sensitivities of CHO V3 cells and V3 cells complemented with WT, T4102A, or T4102D. Cells were irradiated at the indicated doses and plated for analysis of survival and colony-forming ability. **(D-E)** Blocking T4102 phosphorylation increases IR-induced chromosome and chromatin breaks (**D**) and chromosome fusions (**E**). Metaphase spreads were performed following treatment with 2 Gy of IR in CHO V3 cells and V3 cells stably expressing YFP-tagged DNA-PK_cs_ WT, T4102A, and T4102D. Chromosome and chromatin breaks as well as inter/intra-chromosome fusions were enumerated in at least 50 cells. Data are presented as mean±SD and significance was assessed via a two-sided student’s t-test. n.s., not significant; ****, p<0.0001.

## Discussion

DNA-PK_cs_ is heavily phosphorylated in response to DSBs (35). The best characterized DNA-PK_cs_ phosphorylation sites include T2609, S2612, T2620, S2624, T2638, and T2647 (collectively called the T2609 or ABCDE cluster) and S2023, S2029, S2041, S2051, and S2056 (termed the S2056 or PQR cluster) and phosphorylation of both clusters is required for NHEJ and modulation of DSB repair pathway choice. The S2056/PQR and T2609/ABCDE phosphorylation clusters have opposing roles in protecting DSBs, with S2056/PQR protecting the DNA ends and T2609/ABCDE promoting DNA end processing (9). Phosphorylation of the T2609/ABCDE cluster functions by destabilizing the binding of DNA-PK_cs_ to the DNA-Ku complex, triggering its release from DSBs to allow access of DNA ends for processing, including the terminal ligation step of NHEJ (36). The kinase activity of DNA-PK_cs_ is modulated by phosphorylation at T3950 and S56/S72, whereas phosphorylation at T946/S1004 inhibits NHEJ without affecting the enzymatic activity of DNA-PK_cs_ (37). Here, we add to the importance of DNA-PK_cs_ phosphorylation in modulating its functionality, as we show that in response to DSBs, ATM phosphorylates DNA-PK_cs_ in the FATC domain of the protein at T4102. This phosphorylation event promotes stabilization of the DNA-PK_cs_-Ku interaction and the stabilization of the NHEJ machinery at DSBs, both of which stimulate NHEJ.

A body of evidence shows that crosstalk occurs between DNA-PK_cs_ and ATM, and that they may cooperatively initiate DSB repair signaling and regulate DSB repair (20,31). First, combined deficiency of DNA-PK_cs_ and ATM leads to synthetic lethality in mice (38,39). Second, DNA-PK_cs_ and ATM phosphorylate many common targets required for DDR signaling and DSB repair, including H2AX, KAP1, and components of the NHEJ pathway, such as XLF and Artemis (2). Third, DNA-PK_cs_ and ATM phosphorylate each other to modulate specific activities. For example, ATM phosphorylates DNA-PK_cs_ at S3205 and at the T2609/ABCDE cluster in response to DNA damage (37). ATM-mediated phosphorylation of the T2609/ABCDE cluster is essential for NHEJ and allows the freeing of DNA ends for processing by factors including the endonuclease Artemis and the dissociation of DNA-PK_cs_ to allow DNA end ligation (40–42). Moreover, ATM phosphorylation of DNA-PK_cs_ overcomes DNA-PK_cs_-Ku inhibition of resection in vitro (43). Lastly, DNA-PK_cs_ inhibits ATM activity upon DNA damage via phosphorylation at multiple sites to regulate ATM-mediated signaling (44). This study adds to the interplay between DNA-PK_cs_ and ATM and supports another positive role in ATM promoting NHEJ via phosphorylation of DNA-PK_cs_.

A significant number of single molecule studies have produced data showing the makeup of the NHEJ machinery at DNA ends. In particular, two synaptic complexes, the long-range and short-range complexes, have been identified and these complexes are believed to protect the DSB ends and then position them for ligation, respectively (6,7). Although powerful, these studies provide limited insight into the dynamics of the NHEJ machinery at DSBs or how DNA end processing enzymes are recruited. Here, we provide evidence that phosphorylation of DNA-PK_cs_ in its FATC domain at T4102 promotes stabilization of DNA-PK_cs_ at DSBs, which supports assembly and maintenance of the NHEJ core complex at the DNA damage site. We postulate that T4102 phosphorylation of DNA-PK_cs_ induces a conformational change in the FATC domain that is important for mediating DNA-PK_cs_-mediated protein-protein interactions with the core NHEJ factors. The data does not provide evidence if this is direct with the Ku heterodimer or another of the core NHEJ factors, but we hypothesize that this phosphorylation event likely induces an allosteric conformational change that allows the stabilization of the long-range complex. It is also possible that the decrease in the NHEJ core complex stability at the DNA damage site it due to the attenuated DNA-PK_cs_ kinase activity in the T4102A mutant. Furthermore, it also promotes the recruitment of the DNA end processing factor PNKP, suggesting this conformational change upon DNA-PK_cs_ phosphorylation at T4102 may be required for processing of unligatable DSBs. We postulate that this allows ATM protein to ensure the complete repair of DSBs by stabilizing the NHEJ complexes to favor the processing and correct joining of DNA ends.

## Acknowledgements

We thank Linfeng Chi and Shih-Ya Wang for generation of the T4102A and T4102D point mutations and the initial characterization of the mutations. We thank all members of the Davis lab for helpful discussions. A.J.D. was supported by grants by the National Institute of Health (R01-CA162804 and R01-GM04725). Z.D.N. was supported by grants by the National Institute of Health (P01CA092584, P30ES000002). Z.D.N. and D.J.L were both supported by a grant from the American Cancer Society (RSG-22-038-01-DMC). G.J.W was supported by the Natural Sciences and Engineering Council of Canada (4 Department of Biochemistry and Molecular Biology, Robson DNA Science Centre, Arnie Charbonneau Cancer Institute, Cumming School of Medicine, University of Calgary, Calgary, Canada (RGPIN-2018-04327).

## Author contributions

A.J.D. and H.L. conceptualized the project. H.L., Q.Z., D.J.L, A.P., and A.J.D. performed all of the experiments. B.PC.C generated the phosphorylation-specific antibody for T4102. G.J.W. assisted and generated Figure 1B. Z.D.N. planned and was technical advisor for FM-HCR experiments reported in Figure 4B. A.J.D acquired funding. A.J.D and H.L. co-wrote the manuscript. All authors contributed to revising and editing the manuscript.

## Data availability

All study materials will be made available to other researchers. Please contact Anthony J. Davis for reagents.

## Supplementary data

Supplementary Data are available online.

