## Supplemental figures for "ATM phosphorylates the FATC domain of DNA-PK_cs_ at threonine 4102 to promote non-homologous end joining"

### Figure S1

| Accession number | Sequence | Species |
|---|---|---|
| **Mammals** | | |
| hsa_5591 | LSEETQVKCLMDQATDPNILGRTWEGWEPWM | Homo sapiens (human) |
| pps_100981342 | LSEETQVKCLMDQATDPNILGRTWEGWEPWM | Pan paniscus (bonobo) |
| ggo_101128261 | LSEETQVKCLMDQATDPNILGRTWEGWEPWM | Gorilla gorilla gorilla (western lowland gorilla) |
| nle_100586225 | LSEEIQVKCLMDQATDPNILGRTWEGWEPWM | Nomascus leucogenys (northern white-cheeked gibbon) |
| ptr_464165 | LSEETQVKCLMDQATDPNILGRTWEGWEPWM | Pan troglodytes (chimpanzee) |
| pon_100460398 | LSEETQVKCLMDQATDPNILGRTWEGWEPWM | Pongo abelii (Sumatran orangutan) |
| mcc_708029 | LSEETQVKCLIDQATDPNILGRTWEGWEPWM | Macaca mulatta (rhesus monkey) |
| tge_112630683 | LSEETQVKCLIDQATDPNILGRTWEGWEPWM | Theropithecus gelada (gelada) |
| panu_100999716 | LSEETQVKCLIDQATDPNILGRTWEGWEPWM | Papio anubis (olive baboon) |
| mmur_105885240 | LSEETQVRCLIDQATDANILGRTWEGWEPWM | Microcebus murinus (gray mouse lemur) |
| shr_100915132 | LSEETQVKCLMDQATDPNVLGRTWIGWEPWM | Sarcophilus harrisii (Tasmanian devil) |
| cpoc_100724826 | LSVETQVRCLLDQATDPNILGRTWEGWEPWM | Cavia porcellus (domestic guinea pig) |
| pcw_110212506 | LSEETQVKCLMDQATDPNVLGRTWTGWEPWM | Phascolarctos cinereus (koala) |
| cge_100770748 | LSEETQVKCLVDQATDPNILGRTWEGWEPWM | Cricetulus griseus (Chinese hamster) |
| morg_121464862 | LSEETQVKCLVDQATDPNILGRTWEGWEPWI | Microtus oregoni (creeping vole) |
| mmu_19090 | LSEETQVKCLVDQATDPNILGRTWEGWEPWM | Mus musculus (house mouse) |
| rno_360748 | LSEETQVKCLVDQATDPNILGRTWEGWEPWM | Rattus norvegicus (rat) |
| ncar_124981191 | LSEETQVKCLLDQATDPNILGRTWEGWEPWM | Sciurus carolinensis (gray squirrel) |
| opi_101519750 | LSAETQVKCLIDQATDPNVLGRTWAGWEPWM | Ochotona princeps (American pika) |
| **Birds** | | |
| dpub_104303091 | LSEETQVRCLIDQATDPNILGRVWEGWEPWM | Dryobates pubescens (Downy woodpecker) |
| arow_112965430 | LSEETQVKCLIDQATDPNVLGRVWEGWEPWM | Apteryx rowi (Okarito brown kiwi) |
| scam_104149477 | LSEETQVKCLIDQATDPNILGRAWEGWEPWM | Struthio camelus australis (South African ostrich) |
| hald_104317668 | LSEETQVRCLIDQATDPNVLGRVWEGWEPWM | Haliaeetus albicilla (white-tailed eagle) |
| afor_103904238 | LSEETQVRCLIDQATDPNILGRVWEGWEPWM | Aptenodytes forsteri (emperor penguin) |
| **Reptiles** | | |
| amj_102575311 | LTEETQVKCLIDQATDPNILGRVWAGWESWM | Alligator mississippiensis (American alligator) |
| cpoo_109314867 | LTEETQVKCLIDQATDPNILGRVWAGWESWM | Crocodylus porosus (Australian saltwater crocodile) |
| pvt_110077653 | LSEETQVKCLIDQATDPNILGRVWEGWEPWM | Pogona vitticeps (central bearded dragon): |
| sund_121929123 | LSEETQVKCLIDQATDPNILGRVWEGWEPWI | Sceloporus undulatus (fence lizard) |
| **Amphibians** | | |
| xla_373602 | LTEETQVQCLIDQATDPNILGRVWKGWEPWI | Xenopus laevis (African clawed frog) |
| npr_108792226 | LSEETQVQCLIDQATDPNILGRAWKGWEPWI | Nanorana parkeri (Xizang Plateau frog) |
| XP_029447109.1 | LTEETQVKCLLDQATDPNILGRAWQQWEPWM | Rhinatrema bivittatum (two-lined caecilian) |
| **Fish** | | |
| Lcm_102352949 | LPVETQVACLIDQATDPNILGRVWEGWEPWM | Latimeria chalumnae (coelacanth) |
| Arut_117435350 | LSVETQVECLIDQATDPNILGRVWVGWESWV | Acipenser ruthenus (sterlet) |
| Pspa_121314412 | LSVETQVECLIDQATDPNILGRVWAGWEPWV | Polyodon spathula (Mississippi paddlefish) |

**Figure S1. Related to Fig. 1. Conservation of amino acid threonine 4102 (T4102) in DNA-PK$_{cs}$ in vertebrates.** Amino acid alignment of the FATC domain of DNA-PK$_{cs}$ from a variety of vertebrate species., from human to fish. T4102 is indicated in red.

**Figure S2**

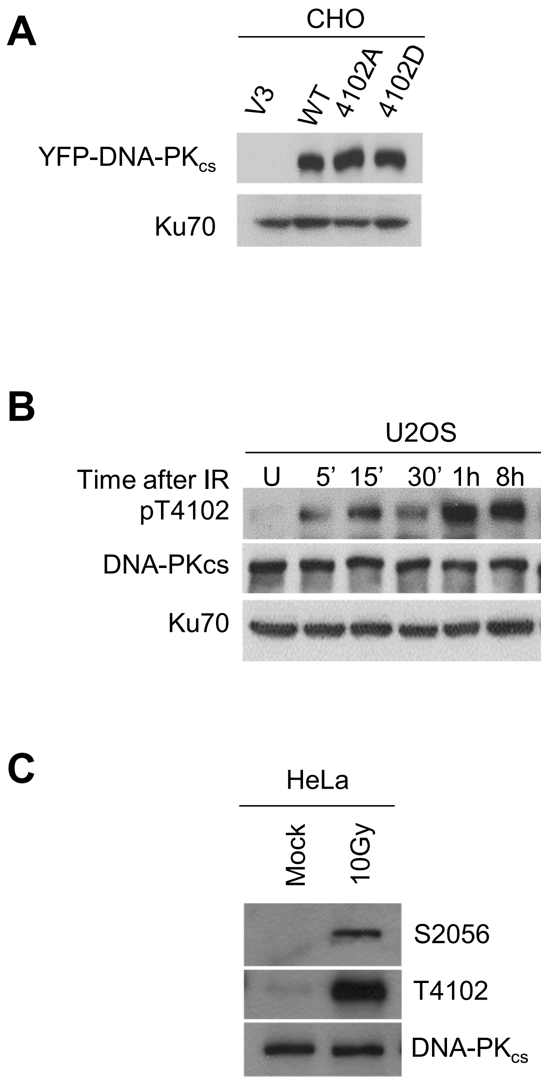


**Figure S2. Related to Fig. 1. DNA-PK$_{cs}$ is phosphorylated at T4102 in response to ionizing radiation (IR) in human cell lines. (A)** Immunoblotting showing expression of DNA-PK$_{cs}$ in CHO V3 cells and V3 cells stably expressing YFP-tagged DNA-PK$_{cs}$ wild type (WT), phosphorylation-null mutant (T4102A), and phosphorylation-mimic mutant (T4102D). **(B)** Phosphorylation of DNA-PK$_{cs}$ at T4102 at the indicated time following exposure to 10 Gy of IR in the human cell line U2OS. **(C)** Phosphorylation of DNA-PK$_{cs}$ at T4102 30 min post-IR (10 Gy) in the human cell line HeLa.

### Figure S3

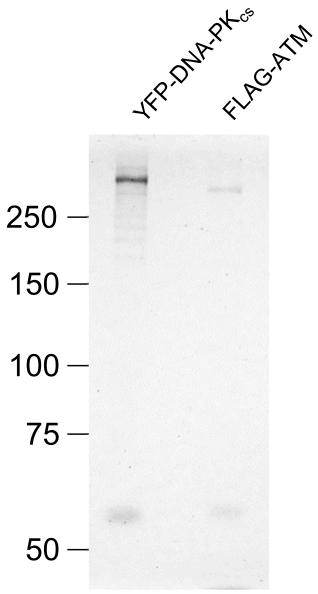


**Figure S3. Related to Fig. 1. Coomassie Blue staining of purified YFP-DNA-PKcs and FLAG-ATM.** YFP-tagged DNA-$PK_{cs}$ was purified from CHO V3 cells stably expressing YFP-DNA-$PK_{cs}$. FLAG-ATM was purified from irradiated HT1080 cells stably expressing FLAG-YFP-ATM.

**Figure S4**

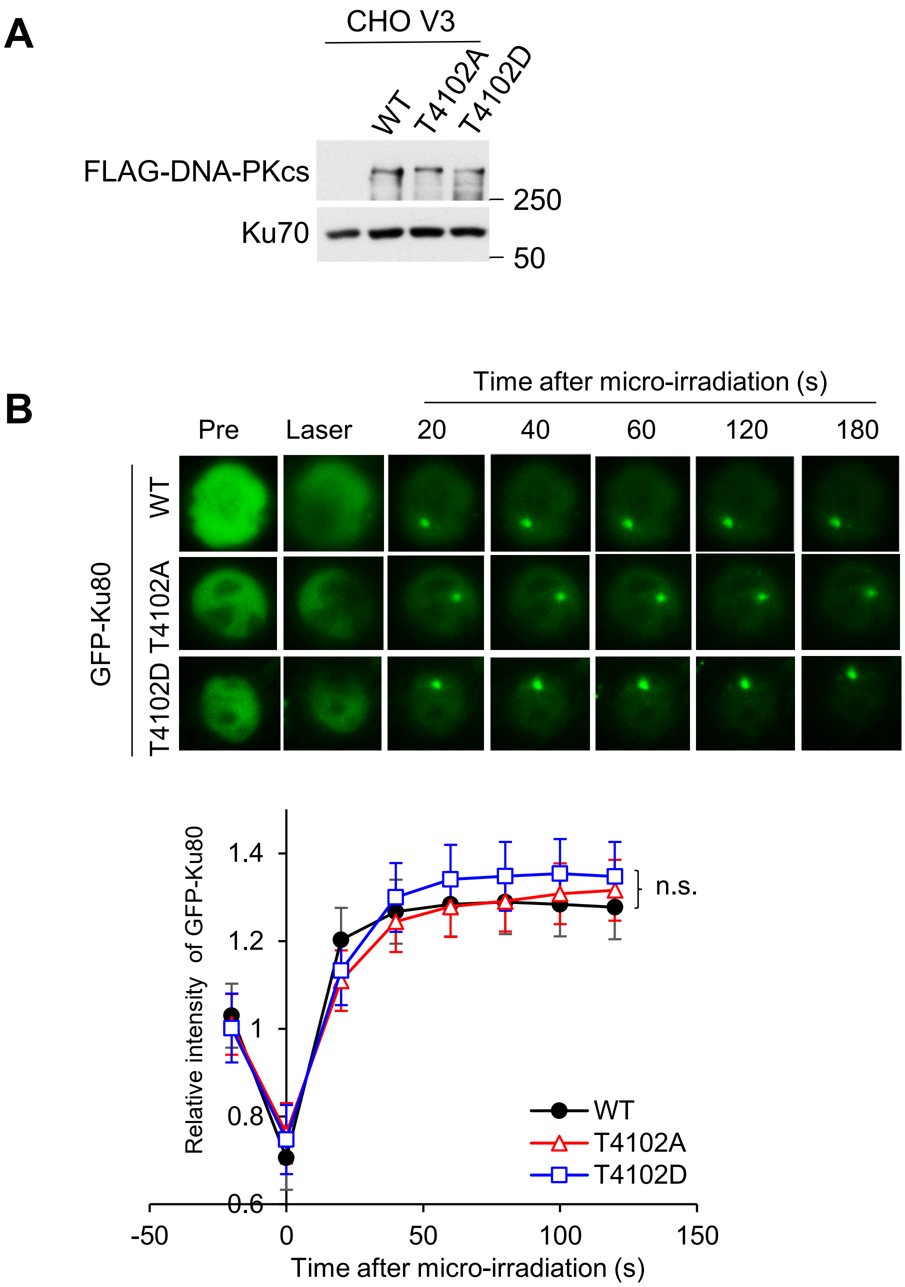


**Figure S4. Related to Fig. 2. Phosphorylation of DNA-PK$_{cs}$ at T4102 is not required for the recruitment of Ku80 to laser-induced DSBs. (A)** Immunoblotting showing expression of DNA-PK$_{cs}$ in CHO V3 cells and V3 cells stably expressing FLAG-tagged DNA-PKcs wild type (WT), phosphorylation-null mutant (T4102A) and phosphorylation-mimic mutant (T4102D). **(B)** Recruitment of GFP-Ku80 was monitored in CHO V3 cells stably expressing FLAG-tagged DNA-PK$_{cs}$ WT, T4102A, and T4102D. The data are presented as mean$\pm$SEM with p-values calculated via student's t-test. n.s., not significant.

**Figure S5**

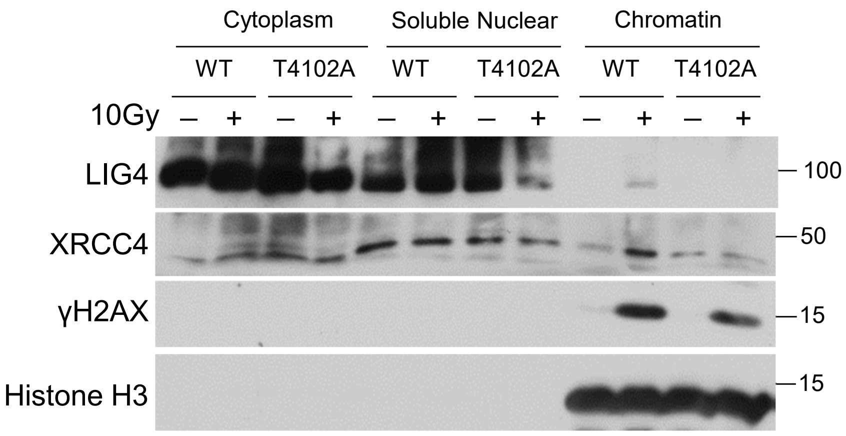


**Figure S5. Related to Fig. 3. Phosphorylation of DNA-PK$_{cs}$ at T4102 promotes the recruitment of the core NHEJ core LIG4 and XRCC4 at to the chromatin fraction following IR.** V3 cells complemented with DNA-PKcs WT or T4102A were mock treated or irradiated with a dose of 10 Gy and allowed to recover for 10 min. Subsequently, cytoplasmic, soluble nuclear, and chromatin fractions were isolated for immunoblotting to examine the recruitment of the proteins listed in the figure to the chromatin fraction after IR.

**Figure S6**

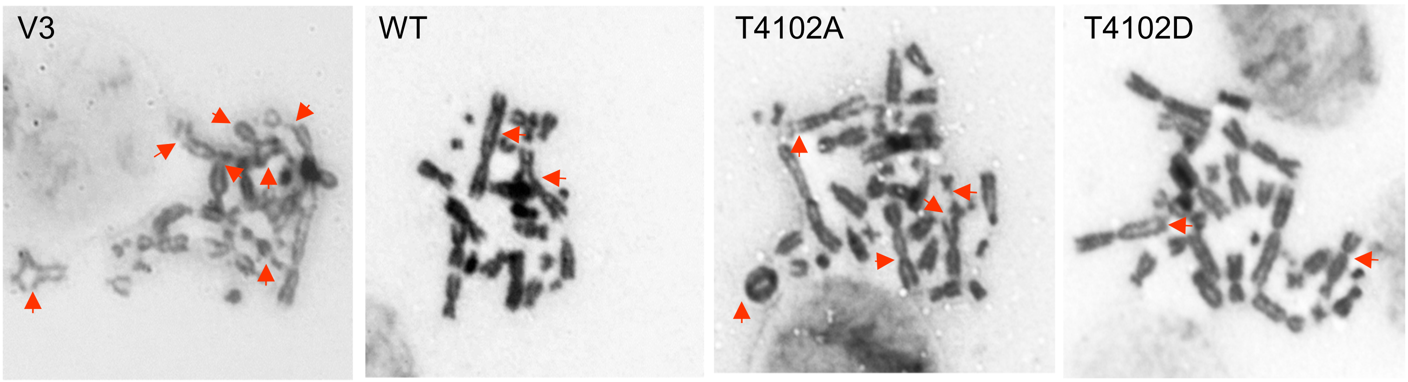


**Figure S6. Related to Fig. 4. Blocking T4102 phosphorylation results in increased IR-induced chromosomal aberrations.** Representative images of metaphase spreads of chromosomes from irradiated CHO V3 cells and V3 cells stably expressing YFP-tagged DNA-PK$_{cs}$ WT, T4102A, and T4102D. Abnormal chromosome/chromatin are labeled with red arrows.
